# Cigarette smoke impairs the endocytotic process in *Saccharomyces cerevisiae*

**DOI:** 10.1101/2024.10.07.616931

**Authors:** Aditya Shukla, Srimonti Sarkar, Alok Kumar Sil

**Author notes:** Corresponding author Alok Kumar Sil.

## Abstract

The accumulation of misfolded proteins inside the cells has been considered to be an important contributor to the development of cigarette smoke-mediated diseases. Since endocytosis plays a crucial role in protein trafficking and clearance, impaired endocytosis may contribute to cigarette smoke-mediated protein accumulation. Therefore, the current study investigated the effects of cigarette smoke extract (CSE) on the endocytosis process in yeast *Saccharomyces cerevisiae*. The current study showed that treatment of cells with CSE caused reduced uptake of FM4-64 stain, indicating impaired endocytosis. Further analysis revealed that CSE treatment resulted in a defect in the recruitment of proteins involved in endocytosis. Also, aberrant actin morphology was found upon CSE treatment, which might interfere with vesicle budding from the membrane. Moreover, the current study showed that the PI4,5P2 level in the plasma membrane in CSE treated cells is reduced due to the failed translocation of MSS4 kinase to the membrane. This reduced PI4,5P2 results in aberrant actin morphology. Thus, the current study demonstrates that CSE treatment causes endocytosis defects and provides insight into this defective endocytosis.

## Introduction

Oxidative stress results from an imbalance between oxidative and antioxidative potentials within the cell. Such an imbalance results due to either an excess or dysregulated generation of prooxidants, and/or a malfunction in the antioxidant system (Ambrose and Barua, 2004). Oxidative stress has been established as a key contributor towards the etiology of cigarette smoking-related illnesses (Boukhenouna et al., 2018, Magder, 2006). Cigarette smoke (CS) is known to contain more than 4700 chemicals, including numerous polycyclic aromatic hydrocarbons and both reactive oxygen and nitrogen species (ROS and RNS). These range from short-lived oxidants like superoxide to long-lived organic radicals like semiquinone [Seo Yoon-Seok et al.,2023]. The presence of these components directly or indirectly causes oxidative stress inside the cell and can affect cellular function by damaging proteins, lipids, nucleic acid, etc. Unlike other oxidative stress-inducing agents, such as H_2_O_2_, CS-mediated oxidative stress acts via a variety of pathways that ultimately result in overloading cells with ROS (Seo Yoon-Seok et al.,2023). It contains oxidants and can also produce oxidants through chemical interactions with biomolecules. In addition, CS can activate cellular ROS sources to increase the production of ROS. Furthermore, CS compromises the antioxidant system, exacerbating the creation of ROS and its effects (Seo Yoon-Seok et al.,2023). Thus, CS is more potent as an oxidizing agent than other oxidizing agents commonly used to test the effect(s) of oxidative stress on different biological events. Consistently CS causes significant harm to cells and increases the risk of either cell death or the occurrence of several diseases such as cancer, COPD, atherosclerosis, etc. (Timothy and Nneli, 2007; Walser et al., 2008; Huang et al., 2019).

One of the key molecular changes induced by CS-mediated oxidative stress is the structural alterations in proteins. This leads to the accumulation of considerable amounts misfolded proteins that leads to proteostasis imbalance. Such imbalance is the underlying cause for pathological conditions like emphysema, cardiovascular diseases, Parkinson’s and Alzheimer’s disease, cancer etc. (Merciniak et al., 2006; Hoozemans et al., 2007; Yoshida, 2007: Tran et al., 2015).

Endocytosis is a fundamental process that plays a crucial role in cellular physiological processes such as protein trafficking to protein clearance (Tofaris et al., 2011; Sahu et al., 2011; Sugeno et al., 2014;uytterhoeven et al 2015;Leibiger et al.,2018). Consistent with this, defects in endocytosis have been implicated in several neurodegenerative diseases (Cataldo et al., 1997; Cataldo et al., 2000; Cataldo et al., 2004). But whether the exposure of cells to CS causes endocytosis defect remains unaddressed. To this end, the present study investigated the effect of CS extract (CSE) on the endocytosis process using *S. cerevisiae*. The usefulness of this model organism for investigating various cellular processes, including endocytosis, stems from the fact that not only is it genetically and biochemically tractable, many of it’s cellular pathways are very similar to that of mammalian cells. In addition, *S. cerevisiae* has already been established as a useful model to investigate the effect of CSE on different aspects of cellular physiology (Biddick and Young, 2009; Das et al., 2014. Maiti et al., 2021).

In yeast, the endocytic process involves a variety of proteins that gather cargo into a membrane patch, invaginate the membrane, and pinch off a vesicle sequentially in various phases (Goode et al., 2015; Boettner et al., 2012; Weinberg and Drubin, 2012; Kirchhausen et al., 2014; Carroll et al., 2012; Brach et al., 2014). While the early-phase proteins (e.g. Ede1, Syp1,End3) have a role in cargo binding and endocytic site location the late-phase proteins (e.g. Sla1,Rvs167,Rvs161) function in promoting membrane invagination and facilitating scission through stimulating actin assembly (Brach et al., 2014, Mooren et al., 2012). On the other hand, some proteins (e.g. Sla2) arrive at the interface between the early and late phases (Kaksonen et al., 2003; Sun et al., 2005; Skruzny et al., 2012), deliver the forces produced by the actin cytoskeleton to the plasma membrane which facilitates the inward motions of endocytic buds. In addition to different proteins, phosphoinositide PI4,5P2 plays crucial role by maintaining actin cytoskeleton organization through interaction with Sla2 (Antonescu et al., 2010; Aguilar and Mas, 2003).

Towards understanding the effect of CSE on endocytosis, the result from this study documented that the treatment of cells with CSE results in defective endocytosis. Further, the results showed that both early and late endocytic markers such as Ede1,End3 and Sla1 are not recruited to the cell membrane upon CSE treatment. The result also showed that CSE treatment causes reduction in the membrane PI4,5P2 level resulting in an impaired recruitment of Sla2 that leads to disorganization of actin filaments.PI4,5P2 Collectively, all these defects result in defective endocytosis in CSE-treated cells.

## Results

### Treatment of cells with CSE affects endocytosis

Cellular uptake of FM4-64 is a well-established method for studying endocytosis as this vital dye is easily incorporated into the plasma membrane and transits via endosomal compartments to the vacuole (Vida et al., 1995). To examine the effect of CSE on endocytosis, *S. cerevisiae* cells were stained with FM4-64 before or after treatment with 15% CSE for different lengths of time. Cells that were stained with FM4-64 after CSE exhibited reduced uptake of dye compared to both CSE-untreated cells and cells that were stained before CSE treatment (**Fig.1A and B**). Cells treated with CSE for 2 h exhibited little or no uptake of FM4-64 (**Fig. 1A**). This result showed that the treatment of cells with CSE caused a defect in the dye uptake, indicating a defect in the endocytosis process.

**Fig. 1.**
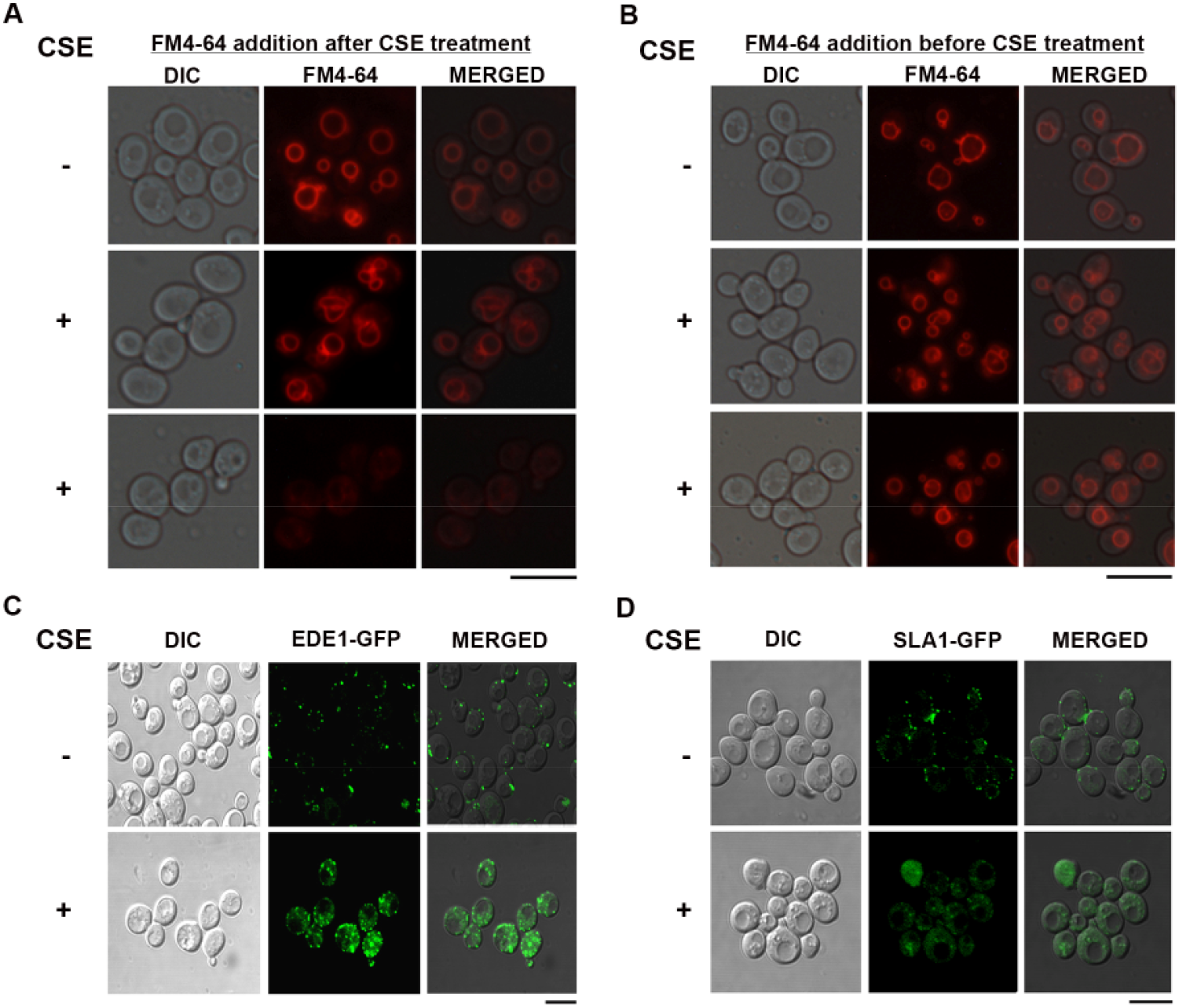
Treatment of cells with CSE affects endocytosis. **A**. Cellular entry of FM4-64 is impaired upon CSE treatment. Exponentially growing *S. cerevisiae* cells were treated with 15% CSE for different periods as indicated and stained with FM4-64 dye. Thereafter. stained cells were visualized under fluorescence microscope. Experiment was performed at least three times independently. Each image is representative of ten different fields. Scale Bar: 10 μm. **B**. CSE treatment after FM4-64 staining does not affect FM4-64 cellular entry. Exponentially growing *S. cerevisiae* cells were stained with FM4-64 and then treated with 15% CSE as indicated. Cells were visualized under fluorescence microscope. Experiment was performed at least three times independently. Each image is representative of ten different fields. Scale Bar: 10 μm. **C, D**. Cells harboring either Ede1-GFP (C), or Sla1-GFP (D) were grown till mid-logphase, treated with CSE (15%) for 2 h, and visualized under a fluorescence microscope. All the experiments were performed at least three times independently. Each image is representative of ten different fields. Scale Bars: 10 μm.

To bolster our observations of endocytosis defect in CSE-treated yeast cells, the sub cellular localizations of Ede1-GFP and Sla1-GFP were examined in CSE-untreated and -treated cells. Ede1 is one of the earliest proteins to reach the nascent endocytic site at the plasma membrane, and it facilitates initiation and maturation of endosomes. On the other hand, Sla1 is a coat protein that arrives to the membrane after Ede1 but before the internalization of the endosome (Weinberg et al., 2012; Goode et al., 2015). Thus, Ede1 and Sla1 constitute ideal markers for the successful recruitment and function of downstream endocytic components. The result showed that in CSE-untreated cells, the punctate GFP signal for these two proteins was very close to the cell surface indicating that they localize to the plasma membrane **(Fig. 1C and D)**. There was little or no signal in the cytosol. In contrast, CSE-treated cells exhibited significant diffused cytosolic signals and a concomitant decrease in the intensity of the signal at the puncta. This indicates a defect in the membrane recruitment of these two proteins, which is likely to impair the initiation of endocytosis.

### CSE affects actin arrangement and tubulin integrity to cause defects in the endocytosis process

The inward movement of the membrane is vital for the formation of endosomes and this is driven by actin-generated force (Marc Abella et al., 2021). It requires polymerization of actin, and therefore it is also possible that CSE impairs actin polymerization. Therefore, the current study investigated actin arrangement in CSE-treated and -untreated cells using phalloidin dye. The results showed that unlike CSE-untreated cells where actin was polarized, clumped or depolarized actin was found in CSE-treated cells (Fig 2A). It indicates that CSE treatment disrupts actin arrangements, and thus it contributes to CSE-mediated endocytosis defect. In addition, the tubulin network also plays a crucial role in endosome movement (Granger et al., 2014). To this end, the current study also examined the effect of CSE on tubulin integrity using Tub3-GFP and found that CSE disrupts the tubulin network (Fig 2B).

**Fig. 2.**
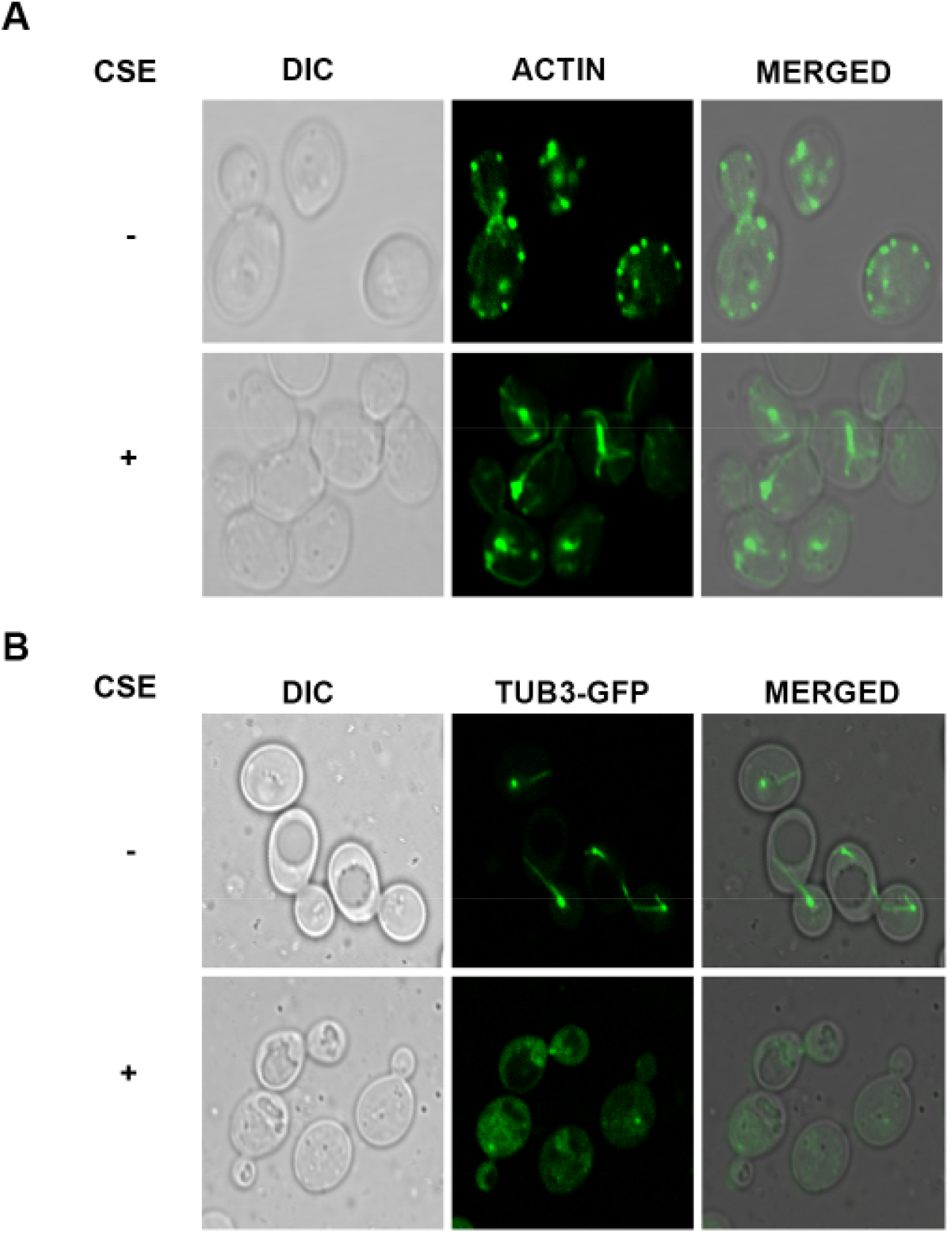
CSE affects actin arrangement and tubulin integrity. **A**. Actin profile. Cells were grown till the mid-log phase, treated with CSE for 2 h and fixed with formaldehyde. After that, cells were stained with phalloidin dye for 45 mins, washed, and visualized under confocal microscope. Each experiment was performed three times independently. Each image is representative of ten different fields. Scale Bars: 10 μm. **B**. CSE disrupts tubulin. Cells harboring Tub3-GFP were grown till the mid-log phase, treated with CSE for 2 h and visualized under a confocal microscope. Experiments were performed three times independently. Each image is representative of ten different fields. Scale Bars: 10 μm

### CSE reduces membrane PI4,5P2 level

Defective actin arrangement may result from impaired recruitment of Sla2 as it stabilizes actin polymerization and helps in the transmission of force generated by actin to the membrane to facilitate inward movement of endosome (Skruzny et al., 2012). Therefore, the current study examined the localization of Sla2-GFP in CSE-treated cells. While in CSE-untreated cells, Sla2-GFP was found only in the plasma membrane as small puncta, in CSE-treated cells, Sla2-GFP was found mostly in the cytosol (**Fig.3A)**. Since Sla2 recruitment depends on the membrane PI4,5P2 (Skruzny et al., 2012), one of the possibility is treatment of CSE causes reduction in PIP2 level at the membrane. In this context, the current study investigated the membrane PI4,5P2 level using PH-PLCδ-GFP as this domain binds specifically to PI4,5P2. Unlike the control cells wherein the PH-PLCδ-GFP signal was restricted to the membrane, the signal in the CSE-treated cells was found in the cytoplasm as scattered punctate structures. This result shows a depletion of PI4,5P2 at the membrane and explains CSE-mediated reduced Sla2 recruitment at the membrane **(Fig.3B)**. As mentioned Sla2 plays a vital role in actin polymerization, this result provides an insight into CSE-mediated defective polymerization of actin (Skruzny et al., 2012). Again the generation of PI4,5P2 depends on the presence of MSS4 kinase at the membrane. Therefore to gain an insight into the reduced PI4,5P2 level at the membrane the current study examined the location of MSS4 kinase in CSE-treated cells using MSS4-GFP. Consistent with our expectation MSS4-GFP was found on the cellular membrane in the control cells, however, in CSE-treated cells, MSS4-GFP failed to get recruited at the membrane resulting in reduced PI4,5P2 level at the membrane **(Fig 3C**).

**Fig. 3.**
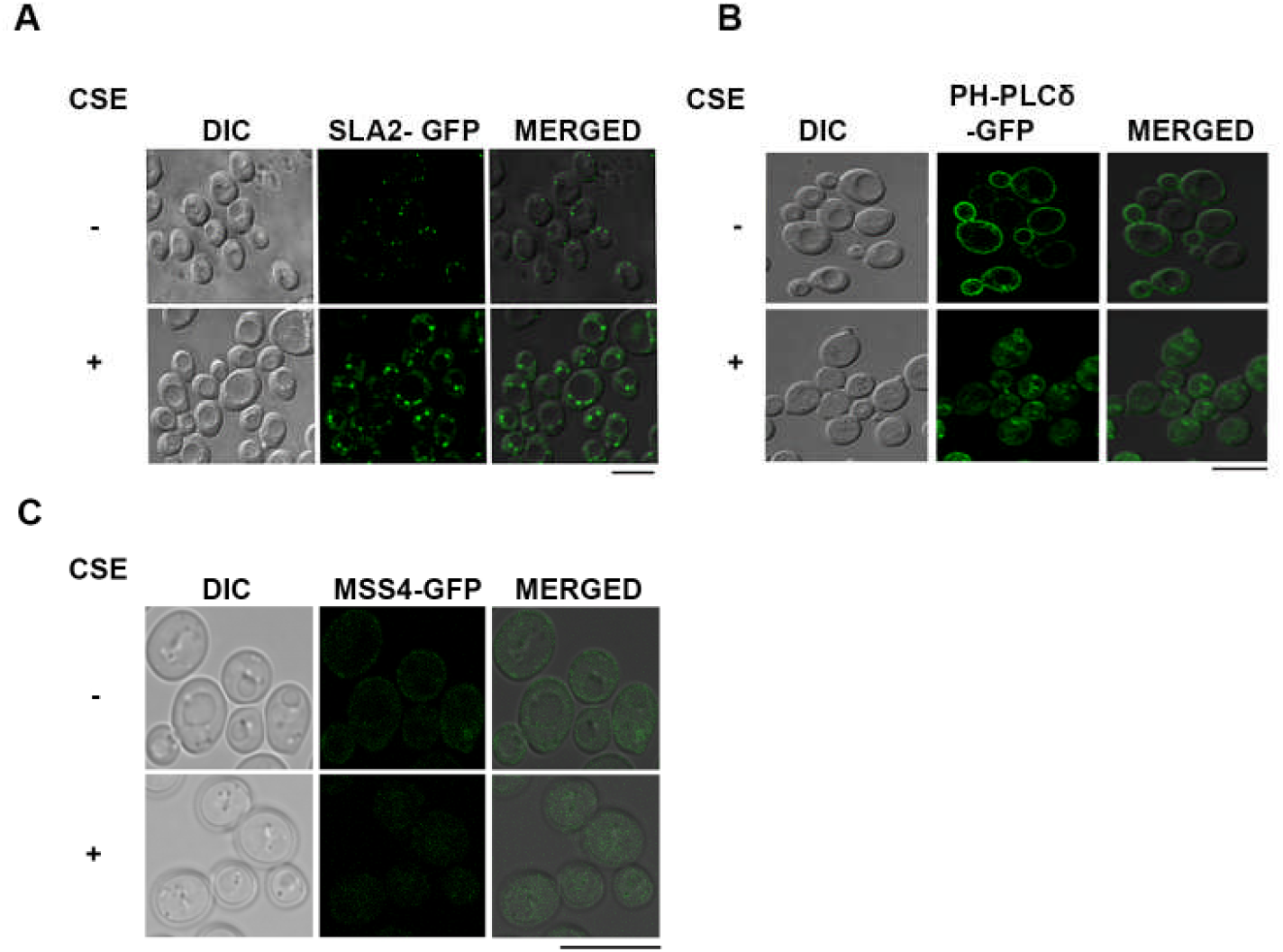
CSE reduces the levels of membrane PI4,5P2. Cells harboring Sla2-GFP (A) or PH-PLCδ-GFP (B) or MSS4-GFP (C) were grown till exponential phase, treated with CSE for 2 h and visualized under confocal microscope. Experiments were performed three times independently, and each image is representative of ten different fields. Scale Bars: 10 μM

### CSE affects receptor recycling

If CSE impairs the endocytosis process, in theory, it should also hinder the recycling of receptors. Therefore to further verify CSE-mediated defective endocytosis, the current study investigated the effect of CSE on the cellular distribution of the Fet3-GFP. Fet3 is a receptor protein and remains at the plasma membrane in resting cells. Again, due to normal turnover, the vacuolar lumen will also be lit up for the presence of GFP in a resting cell. However, if receptor recycling is hampered by defective endocytosis, a part of the Fet3-GFP will be accumulated on the endosomal and vacuolar membranes. Consistent with this, in CSE-untreated control cells Fet3 signal was found on the membrane as well as in the vacuolar lumen **(Fig.4)**, However, in CSE-treated cells, although signal was found on the membrane, a substantial amount of signal was found in the cytosol in the punctate form indicating their localization on the endosome. This result shows defective recycling of Fet3 receptor and thus supports CSE-mediated defective endocytosis.

**Fig. 4.**
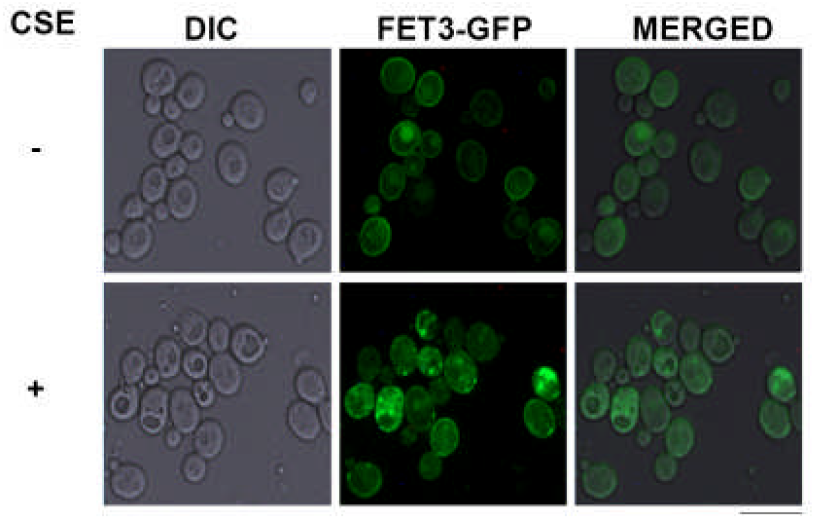
CSE affects Fet3 recycling. CSE treatment causes a defect in Fet3 recycling. *S*.*cerevisiae* cells expressing Fet3-GFP were grown till mid log phase, treated with CSE (15%) for 2 h and visualized under fluorescence microscope. Experiment was performed at least three times independently. Each image is representative of ten different fields. Scale Bar: 10 μm.

## Conclusion

The current study highlights the pleiotropic effects of CSE, which affect different events of endocytosis. Thus, the current study unravels the underlying mechanism of several CSE-mediated diseases, such as Alzheimer’s disease (AD) and Parkinson’s disease (PD), wherein protein accumulation is a significant concern. In addition, previous reports also showed that exposure of cells to CSE impairs nutrient uptake (Das et al., 2014). Based on the results of this study the CSE-mediated defect in nutrient uptake can be attributed to the defective endocytosis as nutrient receptors might fail to internalize or recycle back in the membrane in CSE-treated cells. Moreover, an epidemiology report suggested that during covid pandemic, smokers tend to catch fewer corona virus infections. Again, it can be explained in light of our discovery, as corona virus entry into cells depends on its binding to ACE2 receptor and subsequent internalization via endocytosis. It indicates an endocytotic defect of viral internalization among smokers. In conclusion, the current study has shown how CSE has impacted the process of endocytosis.

## Acknowledgement

The authors would like to acknowledge David G. Drubin and Maya Schuldiner for providing *Saccharomyces cerevisiae* strains used in this study. The authors would also like to acknowledge the Council of Scientific and Industrial Research (CSIR), Govt of India, for providing financial support to AS [09/028(1111)/2019-EMR-1]. The authors would also like to acknowledge the Bose Institute Central Instrumentation Facility and IPLS (University of Calcutta) Instrumentation Facility.

## Author contribution

AS participated in designing experiments, performed all the experiments, analysed the results and participated in writing the manuscript. SS participated in designing experiments, analysed the results and participated in writing the manuscript. AKS conceived the idea, designed experiments, analyzed results, and wrote the manuscript.

## Data availability

All data generated or analyzed during this study are included in this article.

